# Actin reorganization and junctional remodeling in podocyte foot process effacement

**DOI:** 10.64898/2026.11.11.762799

**Authors:** Eva Wiesner, Julia Binz-Lotter, Marie-Kristin Kroll, Katrin Bohl, Jan-Wilm Lackmann, Nelli Rutkowski, Bernhard Schermer, Thomas Benzing, Matthias J. Hackl

**Affiliations:** Department II of Internal Medicine and Center for Molecular Medicine Cologne, University of Cologne, Faculty of Medicine and University Hospital Cologne, 50937 Cologne, North Rhine-Westphalia, Germany; Department of Nephrology, Medical Faculty, University Hospital Düsseldorf, Heinrich-Heine-University Düsseldorf, Düsseldorf, Germany; Cologne Excellence Cluster on Cellular Stress Responses in Age-Associated Diseases (CECAD), University of Cologne, 50931 Cologne, North Rhine-Westphalia, Germany; Center of Excellence in Nephrology, University Hospitals Aachen, Cologne, Dusseldorf, North Rhine-Westphalia, Germany

## Abstract

Damage to podocytes results in kidney disease affecting millions of patients worldwide. Actin remodelling is a shared feature in podocyte foot process (FP) effacement occurring after podocyte injury. However, it remains unclear which processes initiate and orchestrate cytoskeletal changes. Here, we show novel insights into actin dynamics during nephrotoxic serum (NTS) nephritis and other podocyte disease models. Using Lifeact.mScarlet-I reporter expression and STED microscopy in podocytes, we could show distinct actin patterns occurring during FP effacement, independent of damage origin. Additionally, the protein composition of the slit diaphragm changed in a reversible manner. Tight junction proteins ZO-1 and Claudin-5 appeared at the plasma membrane with the onset of proteinuria. The changes in actin and junctional protein assembly during NTS nephritis were reversible as FP effacement disappeared, proteinuria declined and the podocytes recovered. For the first time, we could show with great spatial and temporal detail how the podocyte actin cytoskeleton adapts to insults and that a reversible response in slit diaphragm dynamics leads to a junctional remodelling better suited for upkeeping filtration barrier function.

## Introduction

Podocyte health is essential for the maintenance of the glomerular filtration barrier and overall kidney function. Podocyte injury causes filtration failure, ultimately resulting in proteinuria and elevated serum creatinine levels. Podocytes are specialized epithelial cells with an intricate three-dimensional morphology. Primary processes extend from the cell body and further branch into fine protrusions, the foot processes (FPs). While primary processes predominantly consist of microtubules, FPs rely on actin as their main cytoskeletal component. Adjacent, interdigitating FPs are connected by a specialized cell–cell contact known as the slit diaphragm.

The slit diaphragm contains several podocyte-specific proteins, rendering it a specialized junction. Its protein composition is distinct from that of adherens or tight junctions. Proteins like nephrin and podocin localize to the SD and are essential for its structural and functional integrity. Mutations in their corresponding genes cause hereditary nephrotic syndrome, focal segmental glomerular sclerosis (FSGS) and FP effacement [1], [2]. In addition to unique slit diaphragm proteins, it also contains common tight junction proteins, including ZO-1 and Claudin-5. Their loss results in podocyte injury and proteinuria, highlighting their role in filtration barrier maintenance [3], [4], [5].

Podocytes are non-motile, being permanently anchored to the glomerular basement membrane through integrin-based adhesions. Despite their limited motility, podocytes withstand filtration pressure and shear forces, supposedly through an adaptive, contractile actomyosin network [6], [7]. Disruption of actin dynamics is one feature of podocyte injury. Several mutations in actin-associated genes, like *CD2AP* and *ACTN4*, impair cytoskeletal organization and therefore promote FP effacement [8], [9].

The underlying mechanisms of FP effacement have been studied extensively but are not fully understood yet. It is unclear whether alterations in actin organization initiate downstream signaling for cellular adaptation or whether injury-related signaling drives cytoskeletal remodelling. Most structural insights are based on immunogold-labelled protein electron microscopy. Advancements in imaging techniques enable us to visualize cytoskeletal architecture with more detail. We combined our recently published Lifeact.mScarlet-I mouse model, labelling the actin cytoskeleton in great detail, with a NTS nephritis model, resulting in transient, reversible FP effacement. By studying disease progression at different time points, we aimed to visualize various stages of actin distribution in a temporal context following damage and repair of podocyte FPs.

## Results

### The slit diaphragm can recover after podocyte injury

In order to study actin in FPs effacement, NTS nephritis was induced in 4- to 6-week-old Pod^Lifeact^ mice. This resulted in a transient FP effacement analysed in mice sacrificed on days 2, 3, 5, 9 and 11 after the injection of the serum (Fig.1A). On day 2, the nephrin signal disappeared from the plasma membrane, and FPs were unorganized and effaced (Fig.1B). The nephrin staining for all days showed dots of nephrin in the vicinity of the cell body in circular structures, resembling vesicular internalization (Fig.1B, arrows). At day 2, the NTS model’s heterogeneity led to a wide variation between animals regarding their degree of FP effacement. While some animals still presented FP with a morphology close to control animals, others were severely effaced (Suppl.Fig.1A). This interindividual variation was less pronounced on the following time points, suggesting a higher variability in the disease onset.

**Figure 1:**
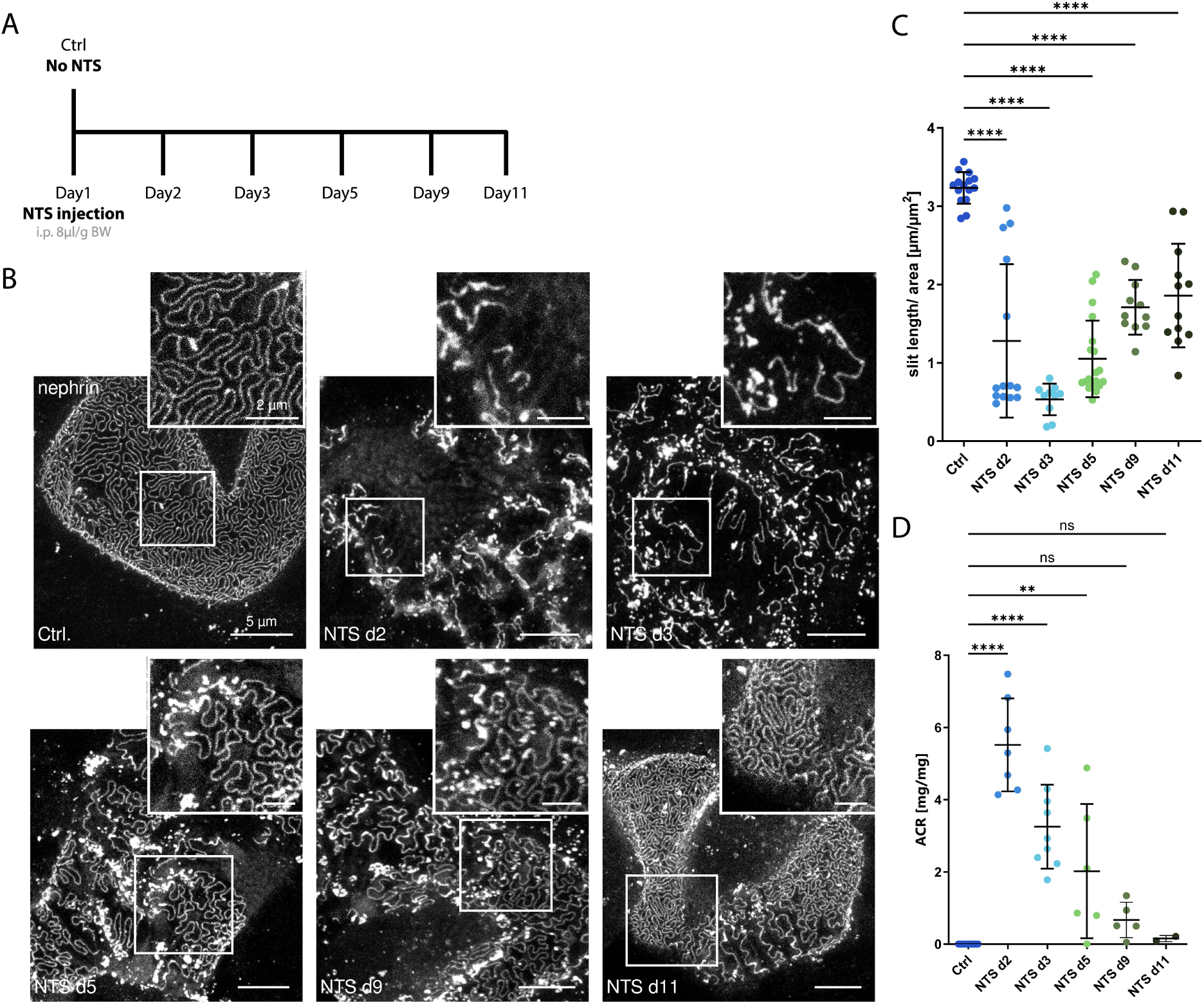
Recovery of the slit diaphragm after NTS induced podocyte injury. A – Mice received a single dose of NTS (8µl/g bodyweight, i.p.). Urine was collected and kidneys were harvested at indicated timepoints after injection. The day of injection was regarded as day 1. B – Representative STED images of anti-nephrin staining of glomerular capillaries shows foot process effacement and internalization of nephrin. C – Ǫuantification of slit diaphragm length/area of images as presented in B using a FIJI macro. D – Albumin creatinine ratio (ACR) of animals injected with NTS. Scale bars – 5 µm, zoom – 2 µm. Statistical test: ordinary one-way ANOVA with multiple comparisons; ns – not significant, p > 0.05, * p ≤ 0.05, ** p ≤ 0.01, *** p ≤ 0.001, **** p ≤ 0.0001. For data points in C: n=15 Ctrl, n=14 NTS d2, n=10 NTS d3, n=19 NTS d5, n=11 NTS d9, n=12 NTS d11; n=glomeruli; For data points in D: n=11 Ctrl, n=7 NTS d2, n=9 NTS d3, n=6 NTS d5, n=5 NTS d9, n=2 NTS d11.

On day 3, the nephrin signal started to increase again, but severe effacement was still present. At day 5 and 9 of nephritis, the slit diaphragm morphology became more regular and elongated, with densely packed FPs and a continuous nephrin staining along the plasma membrane. On day 11, the slit diaphragm pattern recovered, nearly matching control conditions again. FP effacement was still slightly present in some parts of glomeruli, resembling a FSGS-like phenotype. Notably, podocytes’ capacity to reorganize FPs was also highlighted by quantification of the slit diaphragm/area (Fig.1C). Analogous to the slit diaphragm/area, clinical parameters like albumin creatinine ratio (ACR), serum creatinine, and urea peaked already at day 2 of NTS nephritis and recovered to physiological levels by day 11 (Fig.1D, Suppl.Fig.1B and C).

To understand the molecular changes in podocyte FP remodelling after NTS injury, we performed proteomic analysis of glomeruli that were dissected using a laser dissection microscope. We applied the novel approach of assigning a podocyte damage score (PDS) to the samples [10]. The PDS showed a heterogeneous onset of disease on day 2 of NTS nephritis, with the most damage occurring on day 5 and a slight improvement to day 9 and 11 (Suppl.Fig.1D).

### Distinct patterns in actin cytoskeleton are a shared feature of podocyte damage

As FP effacement goes hand in hand with ultrastructural changes, we wondered how it affects the actin cytoskeleton. Therefore, we performed immunofluorescence staining for Lifeact and nephrin in Pod^Lifeact^ mice, thus visualizing the actin cytoskeleton and slit diaphragm pattern simultaneously. In healthy podocytes, actin was mostly present in the FPs, covering the entire capillary surface, but not the podocyte cell body (Fig. 2A). Again, on day 3 of NTS nephritis, the slit diaphragm was mostly depleted of nephrin. The FPs were irregularly shaped and large areas were effaced (Fig. 2B). The Lifeact staining revealed an injury-related actin reorganization that can be sub-categorized into three distinct patterns (Fig.2C). The most frequent pattern (1) showed Lifeact distribution near the cell cortex and at the remains of the slit diaphragm, leaving a ‘empty’ cytoplasmic space with no signal. The second pattern consisted of finger-like actin structures branching from a thick actin fibre resembling former healthy FPs (2). However, they were no longer connected to the membrane or outlined by a nephrin signal. The third pattern was mostly found in severely effaced areas and showed a dotted, irregular Lifeact localization (3). All these changes of the actin cytoskeleton were also reversible, as the Lifeact distribution was similar to controls after 9 and 11 days of NTS nephritis (Suppl.Fig.2A). Figure 2D shows a schematic overview of all three observed patterns.

**Figure 2:**
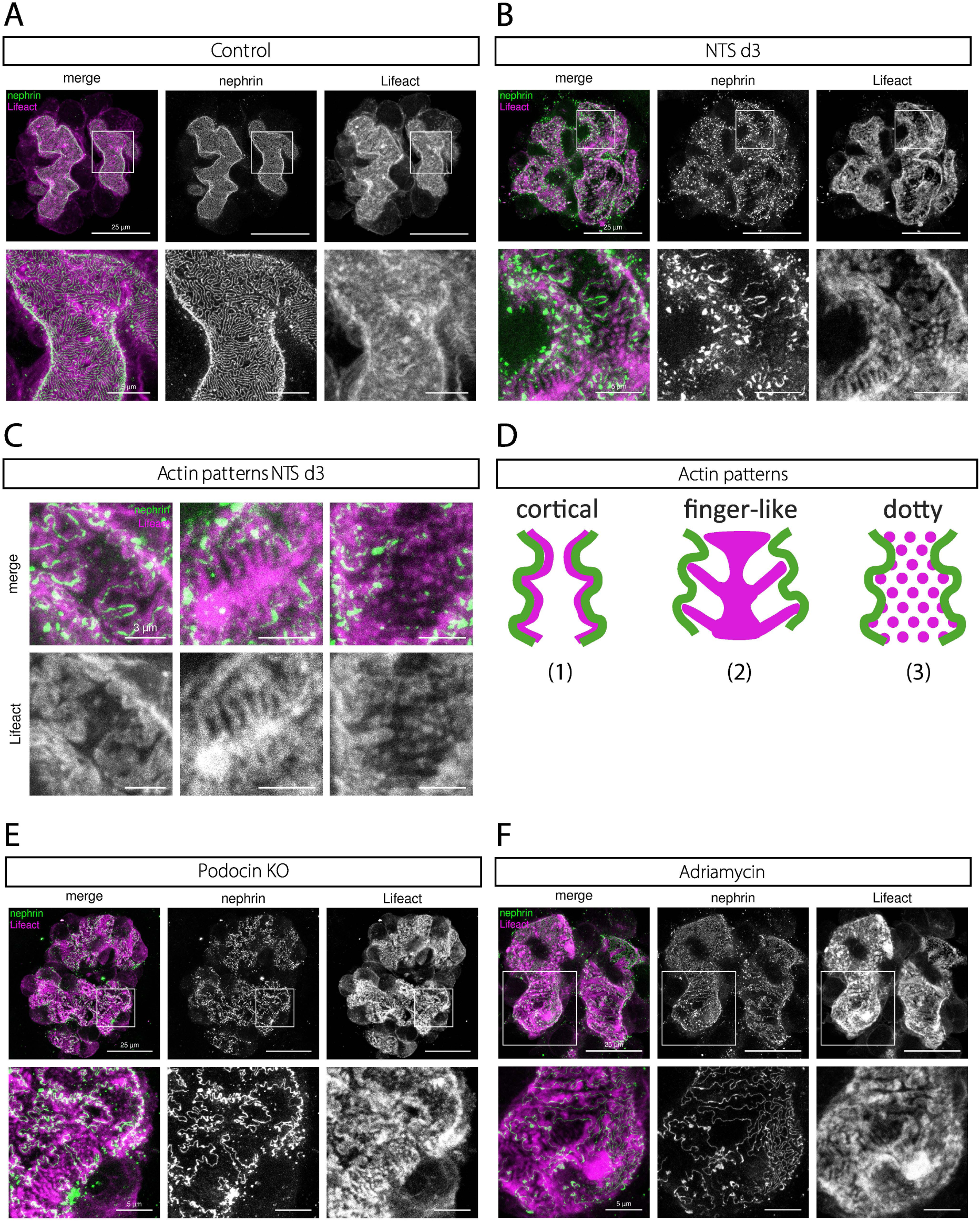
The actin cytosceleton shows distinct patterns upon podocyte injury. A, B, E, F – Representative maximum intensity projection images of glomerular capillary surface stained for nephrin (green) and Lifeact (magenta). Top rows show overview of whole glomerular capillary. Bottom row zoomed in area. A – Healthy control and B – 3 days after a single dose of NTS. C – Representative images of actin patterns found in NTS injected animals (1) ‘cortical’ Lifeact, (2) ‘finger like’ FP like pattern and (3) a ‘dotted’ Lifeact distribution. D – Images of glomerulus of podocin KO mouse. E – Glomerular capillary of adriamycin injected (25mg/g, i.a.) mouse. Scale bars: A,B,E,F: glomeruli – 25 µm, zoom – 5 µm; C – 3 µm.

The endogenous fluorescence of Lifeact.mScarlet-I allowed us to study actin in acute kidney slices (Suppl.Fig.2B). Kidney slices of healthy tamoxifen-inducible iPod^Lifeact^ control animals showed a partial expression of Lifeact.mScarlet-I only in some podocytes and thus enabled a greater contrast in microscopy. Healthy iPod^Lifeact^ mice presented with normal-shaped FPs. NTS-injured animals (NTS d5) showed similar Lifeact patterns as described above. These experiments in vital tissue verified that the observed patterns were not an artifact of sample fixation and processing.

We then wanted to understand if the observed actin patterns are a distinct feature of NTS nephritis or represent a universal pathological change of actin morphology in different podocyte disease models. We generated mice lacking the full-length podocin protein and express the Lifeact reporter in podocytes (Pod^Lifeact podocin KO^). The loss of podocin results in a severe, early onset proteinuria with FP effacement at 4 weeks of age (Fig.2E). We found similar actin patterns as in the NTS model, mostly a cortical (1) and dotty actin distribution (3). Another podocyte disease model used was the adriamycin-induced nephropathy as a chemical injury model (Fig.2F). STED microscopy of nephrin-stained tissue revealed areas with a high degree of effacement, while others were not effaced, a typical hallmark of the resulting FSGS. The observed actin morphology was similar to the two previously shown disease models. Severely effaced regions showed a dotted actin distribution (3). We also observed areas that resembled the cortical actin pattern (1). The finger-like arrangement was mostly missing and could not be seen as frequently as in the NTS nephritis model.

### Actin rearrangements during disease correlate with junctional remodelling

Our data shows a rapid and dynamic reorganization of the actin cytoskeleton in podocytes during NTS nephritis. As the slit diaphragm is a central point in tethering the podocyte plasma membrane to the actin cytoskeleton, we investigated how the junctional composition of the cell-cell junction was affected during FP effacement.

Tight- and adherens junction proteins such as ZO-1, ZO-2, Claudin-5, α-catenin and β-catenin are known to be present in the slit diaphragm, so we performed immunofluorescent labelling of these proteins to investigate their localization during disease onset. In control animals, ZO-1 showed a ubiquitous distribution throughout the podocyte’s FPs (Fig.3A). In injured podocytes, ZO-1 localized mostly to the FP plasma membrane. The other Tjp-family protein, ZO-2, showed only a weak expression in podocytes and its localization was not altered during disease. This was also true for the adherens junction proteins α- and ß-catenin. Surprisingly, the tight junction protein Claudin-5 showed a similar re-localization to the slit diaphragm as ZO-1 in injured podocytes.

**Figure 3:**
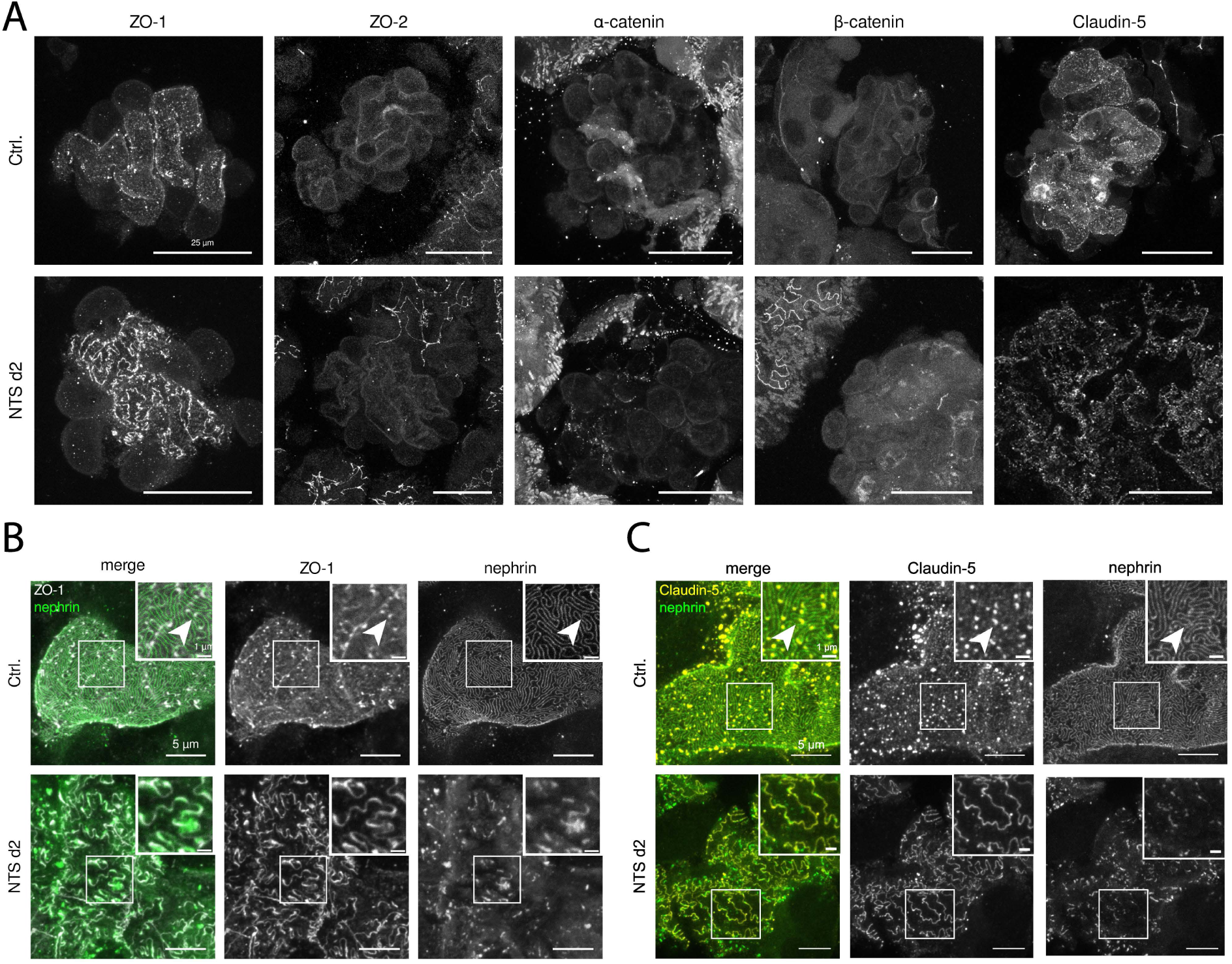
Tight junction proteins re-localize to the SD in FPs effacement. A – Representative images of glomerular capillary surface stained for indicated proteins in healthy control animal (top row) and day 2 of NTS nephritis (bottom). B – Staining for ZO-1 and nephrin in healthy control animal (top row) and day 2 of NTS nephritis (bottom). C – Staining for Claudin-5 and nephrin in healthy control animal (top row) and day 2 of NTS nephritis (bottom). All images are maximum intensity projections. Scale bars: A – 25 µm, B&C – 5 µm, zoom – 1 µm.

High-resolution STED microscopy revealed a detailed view of ZO-1 and Claudin-5 localization in relation to the slit diaphragm during effacement. In healthy animals, ZO-1 could be found throughout the FPs but showed the strongest signal at the distal part of the FP, where membrane curvature is at its maximum (Fig.3B zoom, arrowhead). With disease onset at day 2 of NTS, ZO-1 started to specifically localize to the slit diaphragm. Interestingly, Claudin-5 was found at the same unique spot in FPs as ZO-1 in healthy animals. Upon FP effacement, it was found throughout the whole length of the membrane of the cell-cell junction (Fig.3C).

### ZO-1 re-localization as a hallmark of podocyte effacement

To further investigate ZO-1 re-localization throughout the NTS nephritis course of injury and recovery, we quantified the signal of ZO-1 in relation to the nephrin signal. We utilized a macro-based detection of fluorescent signals. The ratio of detected ZO-1 and nephrin was then normalized to the amount of nephrin detected, giving the percentage difference. This enabled comparisons between time points and disease models. During the time course of NTS nephritis, the percentage difference peaked on day 2 and remained high on day 3 (Fig.4A). Together with the re-expression of nephrin at the slit diaphragm on day 5, the percentage difference declined and reached levels of control animals at day 9 and 11. Representative images of glomeruli from all days stained for Lifeact, ZO-1 and nephrin are shown in Suppl.Fig.2C.

**Figure 4:**
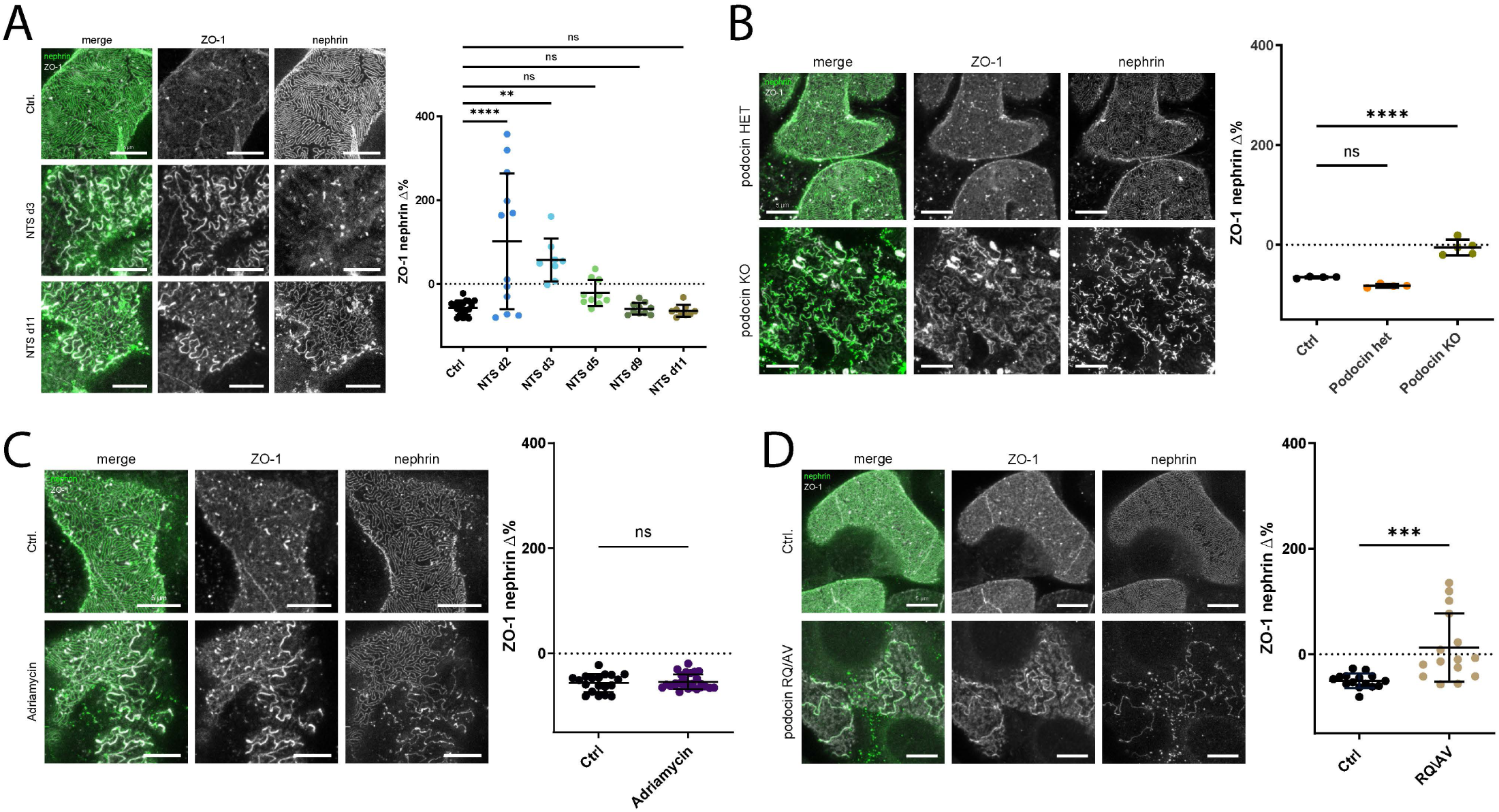
ZO-1 relocates to the slit diaphragm in effaced areas. A-D– Representative maximum intensity projection images of glomerular capillary surface stained for nephrin (green) and ZO-1 (grey). ZO-1 re-localization is quantified by the percentage difference of ZO-1/nephrin signal normalized to the total nephrin signal detected. Images were analyzed using a FIJI macro (s. methods). A – Control and NTS injected animals. Representative images are shown for day 3 and 11 of NTS. B – ZO-1 expression pattern in podocin KO animal (bottom row) and podocin heterozygous control (top row). C – ZO-1 expression pattern in Adriamycin injected (25mg/g, i.a.) mouse (bottom row) and sham surgery control (top row). D – ZO-1 expression pattern in podocin compound heterozygous RǪ/AV animal (bottom row) and control (top row). Scale bars – 5µm. **Data points:** A – n=1G Ctrl, n=12 NTS d2, n=8 NTS d3, n=10 NTS d5, n=10 NTS dG, n=G NTS d11 glomeruli; B – n=4 Ctrl, n=4 podocin Het, n=5 podocin KO; C – n=1G Ctrl, n=27 Adriamycin; D – n=15 Ctrl, n=15 RǪ/AV. Statistical tests: ordinary one-way ANOVA with multiple comparisons for A+B, unpaired t-test for C+D; ns – not significant, p > 0.05, * p ≤ 0.05, ** p ≤ 0.01, *** p ≤ 0.001, **** p ≤ 0.0001.

For the Pod^Lifeact podocin KO^ model, a smaller increase in the percentage deviation of ZO-1/nephrin was observed (Fig.4B). Control animals and animals heterozygous for podocin showed a similar ZO-1/nephrin difference as controls. We also quantified the percentage difference of ZO-1/nephrin in the pharmacological adriamycin disease model (Fig.4C). The average percentage difference did not change compared to sham surgery control animals. In order to exclude the possibility that the occurrence of ZO-1 at the slit diaphragm in effaced areas was influenced by reporter expression, we made use of a genetic injury model: mice carrying a podocin RǪ/AV compound heterozygous mutation but no Lifeact reporter. We observed a similar elevation in ZO-1 and nephrin percentage difference in these compound heterozygous animals to the full _Pod_^Lifeact podocin KO^ _(__Fig.4D__)._

## Discussion

Our work aimed to gain a better understanding of podocyte FP effacement and its effect on actin cytoskeleton remodelling and slit diaphragm junction composition. The NTS nephritis model showed that podocytes have the capacity to recover from immunological injury. Proteinuria levels peaked on day 2 of NTS nephritis and were accompanied by reduced slit diaphragm length and pathological actin patterns. These changes persisted up to day 5 of nephritis. Starting on this day, a more regular, healthy slit diaphragm morphology was reinstated. On days 9 and 11, podocytes showed a mix of recovered areas and effacement, resembling a FSGS phenotype. Despite severe injury, podocytes could recover and reinstate their unique morphology. It remains unclear whether a further, complete repair of FP morphology occurs in even later timepoints. Glomerular crescents, scarring, and filtration loss are known NTS effects after the initial phase of inflammation [11]. Future experiments could help to delineate whether glomeruli progress into forming crescent or completely recover in the setting of our NTS model.

The involvement of actin deregulation in podocyte injury has been known for a long time [12], [13], [14]. Targeting extracellular proteins through the NTS model led to a disruption of the intracellular actin cytoskeleton, most likely by reduction of adhesion points to membrane proteins, especially the slit diaphragm protein complex [15], [16]. Previous research also described an actin mat upon effacement [14]. The actin patterns we observed were well-conserved between different disease causes. Immunological, chemical, and genetic injury caused podocyte damage, leading to distinct patterns, suggesting their occurrence to be at least partially independent of the type of injury.

The cortical, empty actin pattern and the finger-like pattern seem to be mostly present in areas that are not completely effaced. One could hypothesize that it reinforces the filtration barrier by recruiting proteins to the remains of the slit diaphragm. Additionally, the changed actin localisation could reduce the compression of the FP network on the glomerular basement membrane, which could explain the appearance of the proteinuria peak early during the course of disease (NTS day 2) [17]. The appearance of a subcortical actin network is a typical feature of the calcium-mediated actin reset (CaAR) [18]. The formation of the cortical actin pattern could involve calcium-mediated signaling, but its exact role needs to be studied further. The dotty pattern mostly occurred in areas where individual FPs were completely lost and only one flat podocyte cytoplasm covered the capillary. These dotty structures might be anchored to integrins, helping podocytes to attach more firmly to the basement membrane and withstand increased filtration pressure due to effacement [19].

The downregulation of nephrin following NTS injury is likely to mediate a remodelling of the cell-cell contacts between podocytes. The reoccurrence of tight junction proteins during injury, replacing slit diaphragm proteins, has been demonstrated [5], [20], [21], [22]. Through advancements in microscopy, we were able to examine immunolabeled proteins of the slit diaphragm with high resolution, providing more information than electron microscopy. ZO-1 is a component of the slit diaphragm, interacting with NEPH1, Claudin-5, and actin [3], [23], [24], [25]. During kidney development, it is present in podocytes at the early capillary loop stage [26]. Immunofluorescence images revealed higher staining intensity for ZO-1 than for catenins or ZO-2. Upon injury, both ZO-1 and Claudin-5 relocate completely to the cell membrane, shifting the junctional composition towards a tight junction. This might be a form of emergency repair mechanism to prevent proteins from passing the filtration barrier [24], [27]. This hypothesis is also supported by the fact that ZO-1 or Claudin-5 knockout in podocytes causes proteinuria and effacement in mice [3], [4]. As those lines were unconditional knock-out lines, it remains unclear if loss during embryonic development or in mature podocytes is responsible for this phenotype. Further experiments with inducible knockout mouse lines of ZO-1 and Claudin-5 are warranted to delineate the role of these proteins after the slit diaphragm has completely developed.

The special localization of Claudin-5 and ZO-1 at the furthest tip of FPs hints at a special function of this area. The slit diaphragm has been suggested to have properties of tricellular junctions, but further validation is needed to confirm involvement of ILDR2 or tricellulin in the re-localization of tight junction proteins in FPs [28].

The reversible nature of the junctional re-composition highlights how important it is that post-mitotic podocytes react with an adaptive response to injury. It remains unclear what triggers the change in ZO-1 localization upon podocyte damage. We hypothesized that there must be additional signals, and not only the depletion of nephrin or podocin, for tight junction proteins to reinforce the slit diaphragm cell-cell contact. ZO-1 is very prominently recruited to the slit diaphragm in the podocin KO animal, where nephrin is still present and ZO-1 redistribution also occurred in podocin RǪ/AV heterozygous mice.

The correlation of changes in actin patterns and ZO-1 distribution suggests a potential mechanistic link between cytoskeletal properties and the slit diaphragm junction. The actin cytoskeleton is most likely not a direct target of NTS auto-antibodies but was still affected possibly due to changes in molecular tension. ZO-1 conformation has been proposed to function as a molecular tension sensor regulating epithelial homeostasis, depending on the organization of the actin cytoskeleton and myosin-II activity [29]. ZO-1 conformational changes may recruit other proteins to the podocyte membrane, compensating for the loss of podocyte-specific slit diaphragm proteins. The changes might also be triggered by phosphorylation of NEPH1 as a known interactor of ZO-1 or nephrin, with its function as a signaling hub [25], [30], [31], [32], [33].

Our data shows the capacity of podocytes to rebound after damage. However, it was collected for different disease days in different animals. Future work could focus on a longitudinal study design, especially using an abdominal window implanted above the kidney to track changes over time [34]. In conclusion, our data provides novel insights into the reversible nature of FP effacement and how junctional composition and actin dynamics are altered in disease. Podocytes are capable of recovering their slit diaphragm morphology and function upon injury. A universal, adaptive response of the actin cytoskeleton and junctional proteins to FP effacement might provide a future therapeutic target by reinforcing signals for repair and reinstating a functional filtration barrier.

## Materials & Methods

### List of Antibodies

| **Antigen** | **Company & No** | **Dilution** | **RRID** |
|---|---|---|---|
| Nephrin | R&D Systems, AF4269 | 1:100 |  |
| Nephrin | Fitzgerald, 20R-NP002 | 1:100 |  |
| podocin | Sigma, P0372 | 1:100 |  |
| ZO-1 | Invitrogen, 33-9100 | 1:100 | AB_2533147 |
| ZO-2 | Invitrogen, 38-9100 | 1:100 | AB_2533390 |
| RFP | Rockland, 600-401-379 | 1:200 |  |
| Claudin-5 | Invitrogen, 35-2500 | 1:100 | AB_2533200 |
| α-catenin | Sigma, C2081 | 1:100 |  |
| β-catenin | BD Transduction Labs, 610154 | 1:100 | AB_397555 |

### Mice

Animals were derived and maintained in the CECAD Research Center, University of Cologne, Germany, in individually ventilated cages (Greenline GM500; Tecniplast) at 22°C (± 2°C) and a relative humidity of 55% (± 5%) under a 12-hour light cycle on sterilized bedding (Aspen wood, Abedd, Germany). Mice had access to a sterilized commercial pelleted diet (Ssniff Spezialdiäten GmbH) and acidified water ad libitum. The microbiological status was examined as recommended by the Federation of European Laboratory Animal Science Associations (FELASA) and mice were free of all listed agents, including opportunists. Mice (C57BL/6N) of both sexes at the age of 4-6 weeks were used. Mice homozygous for Lifeact.mScarlet-I^fl/fl^ were crossed to GCaMP3^fl/fl^ podocin-Cre^tg/wt^ to achieve Lifeact.mScarlet-I/GCaMP3 heterozygosity in all podocytes (Pod^Lifeact^) [35], [36], [37]. Inducible expression was obtained by using the tamoxifen-responsive podocin-iCreERT2 to generate iPod^Lifeact^ mice [38]. 4-6 week old iPod^Lifeact^ mice received a single Tamoxifen dose of 0.05 mg/g bodyweight via oral gavage. Lifeact.mScarlet-I^fl/fl^ podocin-Cre^tg/wt^ mice were crossed with podocin^fl/fl^ animals to achieve full podocin KO and Lifeact expression in podocytes [39]. Breeding of this mouse line resulted in only one offspring animal of the desired genotype. Due to the severity of the podocin knockout alone, additional Lifeact expression might negatively affect podocytes and further burden renal function, leading to early lethality. Compound heterozygous podocin RǪ/AV animals were available in-house as previously published [17]. No animals were excluded; no randomization was performed. Potential confounders were not controlled and blinding of the study was not performed. N equals the number of mice if not stated otherwise. All experiments were approved by the LAVE NRW, Germany, with the authorization number: 81-02.04.2018.A351 and 81-02.04.2023.A301. Animal experiments complied with LAVE regulations and were conducted in accordance with ARRIVE guidelines.

### Induction of nephrotoxic serum nephritis

Mice received a single dose of nephrotoxic serum (Probetex, LOT #612-3T; 8µl/g bodyweight) as an i.p. injection when 4-6 weeks old. Day 1 is regarded as the day of injection. Mice were euthanized with cervical dislocation on the indicated time points (day 2, 3, 5, 9 and 11). The kidneys were removed, decapsulated and further processed for imaging.

### Induction of Adriamycin induced FSGS

Due to the young age (4-6 weeks) of mice, Adriamycin was administered intravenously with a small surgical procedure. Before surgery, mice received 0.1 mg/kg BW Buprenorphine. Animals were then anesthetized with 5 % isoflurane and the anesthesia was maintained at 1.5%. After a small incision at the neck, the jugular vein was tied cranially and then canulated. Adriamycin (or saline for sham surgery) was slowly injected into the vein with a dose of 25 mg/kg bodyweight. After the injection, the vein was ligated and the skin was closed. Animals were screened for the development of proteinuria and euthanized by cervical dislocation 5 days post-surgery. The kidneys were removed, decapsulated and further processed for imaging.

### Analysis of proteinuria and serum parameters

For the determination of the albumin-to-creatinine ratio (ACR), both albumin and creatinine concentrations were measured in spot urine samples. Urinary albumin levels were quantified by ELISA. Briefly, 96-well plates were coated with an anti–mouse albumin antibody (1:10.000; A90-134A, Bethyl Laboratories) for 1 h at room temperature. Standards and urine samples were added to the plates and incubated for 1 h. Detection was performed using an HRP-conjugated goat anti–mouse albumin antibody (1:25000; A90-134P, Bethyl Laboratories). Absorbance was measured at 450 nm using a microplate reader (Tecan). Urinary creatinine concentrations were determined using a commercially available colorimetric assay kit (Creatinine (Urinary) Colorimetric Assay Kit, 500701, Cayman Chemical). Serum parameters were determined through the Clinical Chemistry of the University Hospital Cologne and measured on a cobas c systems. Creatinine was determined using CREP2 Creatinine plus ver.2 test (Roche). Urea was determined using UREAL test kit (Roche).

### Preparation and imaging of acute kidney slices

Acute slices were prepared according to previously published protocols [40]. Briefly, kidneys were embedded in 4% low-melting agarose and sectioned into 300 μm slices using a vibratome (VT1200 S, Leica). Slices were maintained in carbogenated Krebs–Henseleit buffer (95% O₂/5% CO₂). For imaging, individual slices were transferred to an imaging dish and stabilized with a slice anchor (Warner Instruments) to minimize tissue movement. Confocal imaging was conducted on a STELLARIS 8 STED microscope using an 86× water-immersion objective (HC PL APO 86x/1.2 W motCORR STED W).

### STED sample preparation and imaging

Immunofluorescent labeling was performed as previously described [41]. Kidneys were fixed overnight in 4% paraformaldehyde (PFA) and subsequently incubated in a hydrogel solution for 24h. The hydrogel was polymerized for 3h at 37 °C. After washing, the tissue was embedded in 3% agarose and sectioned into 200 μm slices using a vibratome (VT1200 S, Leica). Sections were optically cleared overnight (200 mM boric acid, 4% SDS, pH 8.5) and then incubated with primary antibodies against the target proteins (see ‘List of antibodies’). Confocal and STED imaging of slices was carried out using a Leica TCS SP8 gSTED 3X microscope equipped with a white light laser (WLL) and a 775 nm STED depletion laser (Leica Microsystems).

### Image processing

Images were processed using FIJI and Omero.figure web application. slit diaphragm length/are analysis and normalized ZO-1/nephrin ratio were determined using a FIJI macro [17]. Figures were assembled using Adobe Illustrator (Version 29.7.1).

### Statistical analysis

For statistical analysis of data, the software Graph Pad Prism (Version 10.6.1. (892) Sep. 8, 2025) was used. We performed an ordinary one-way ANOVA with multiple comparisons or unpaired t-test where necessary. ns – not significant, p > 0.05, * p ≤ 0.05, ** p ≤ 0.01, *** p ≤ 0.001, **** p ≤ 0.0001.

### Laser dissection microscope

For the isolation of glomeruli from mouse kidney tissue, formalin-fixed paraffin-embedded (FFPE) kidney samples were cut into 5 µm sections and mounted on PEN membrane glass slides (Applied Biosystems). Sections were deparaffinized using a brief descending ethanol series and air-dried overnight. The LMD7 Laser Microdissection Microscope by Leica (DFG-INST 216/1179-1 FUGG) was used to cut glomerular sections from kidney tissue. Isolates were collected in mass-spectrometry buffer (20 ml MS-grade water, 1000 mg SDS, 80 mg Tris). Fifteen glomerular sections were isolated per sample. Glomerular isolates were incubated for 10 minutes at 95 °C followed by Bioruptor sonication for 10 cycles with 30/30-s on/off settings. Samples were incubated with dithiothreitol (10 mM) at 55°C for 30 minutes, followed by the addition of chloroacetamide ( 40 mM).

### Proteomics analysis

Proteomic analysis was conducted by the CECAD Proteomics Facility. Samples were digested using a modified SP3 approach [42], foregoing the second day peptide cleanup and instead loading the complete digest on Evotips (Evosep) after acidification following the vendor protocol. Samples were analyzed on an Orbitrap Exploris 480 mass spectrometer (Thermo Scientific, funded by the Deutsche Forschungsgemeinschaft INST 216/1163-1 FUGG) coupled to an Evosep ONE. The Evosep was run with its Whisper Zoom 20 SPD gradient using an Aurora Elite column (Ionopticks). The mass spectrometer was operated using a WHISH-DIA approach [43]. MS1 spectra were acquired between 400 and 1000 m/z with a resolution of 120k, MS2 spectra were acquired in the same range at 60k resolution in 25 m/z windows, resulting in 24 scans total. Fragments were acquired in a range of 250 to 1500 m/z with a normalized AGC target of 1000% and 30% normalized HCD collision energy. Every 6 scans, an MS1 scan was inserted. Samples were analyzed in Spectronaut 19 (Biognosys) after conversion into htrms format using a canonical murine SwissProt database (UP0589, downloaded 15/01/2025). Afterwards, identified and quantified protein groups were exported and analysis was performed on the iBAǪ values. Imputation of missing values was done with R package imputeLCMD using function impute.MinDet with default settings [44]. The resulting data matrix was log transformed for all further analysis. The podocyte damage score (PDS) was calculated on the imputed, log-transformed data matrix using the method “Podocyte Damage Score” and code published in [10] and [45].

## Acknowledgements

We thank S. Keller, M. Brütting, S. Greco-Torres and A. Köser for technical assistance. The authors acknowledge the support of the CECAD Imaging Facility and CECAD Proteomics Facility.

