## Supplemental Information for "Actin reorganization and junctional remodeling in podocyte foot process effacement"

### Supplemental Material

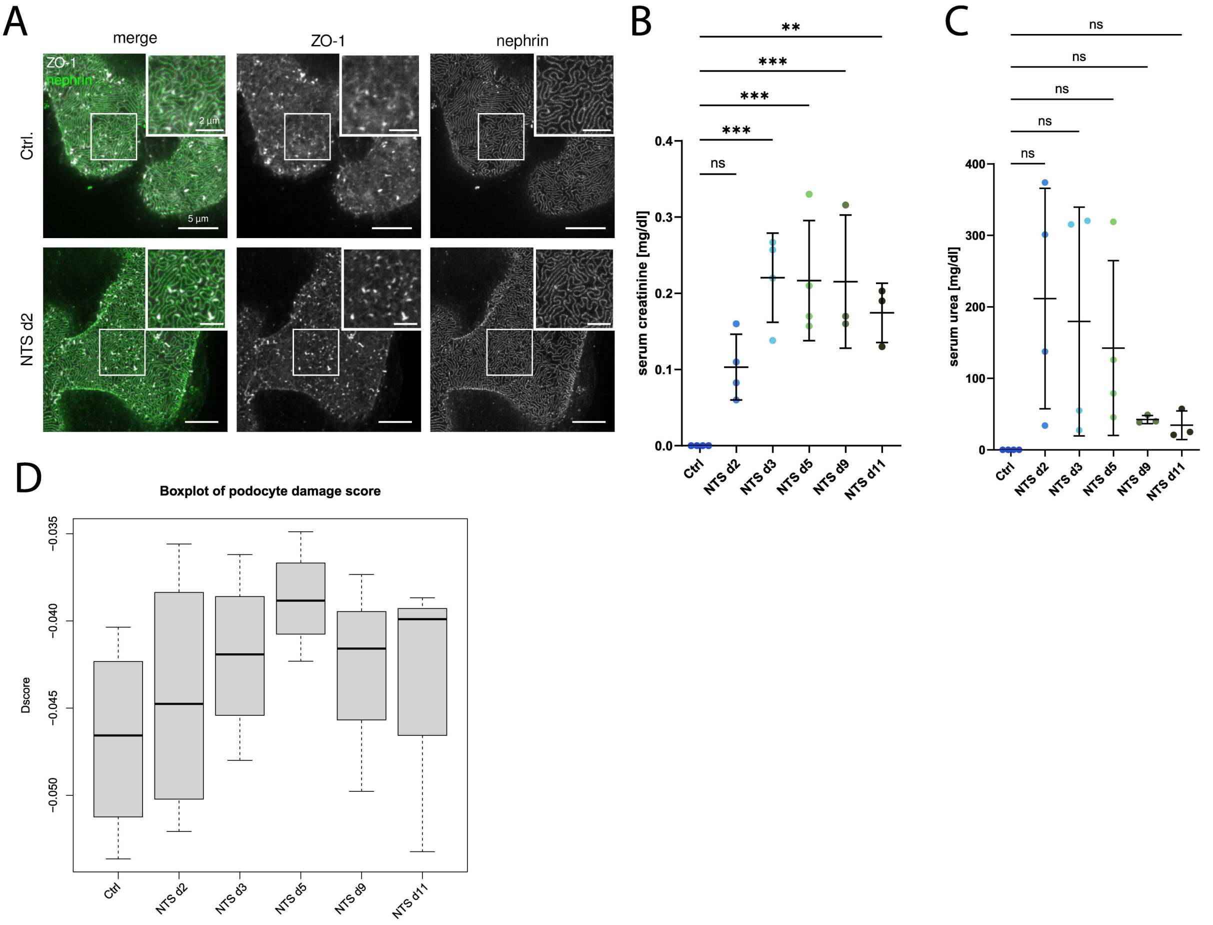


**Supplementary Figure 1: Characterization of NTS induced damage.** A – Representative maximum intensity projections of nephrin (green) and ZO-1 (white) stained mouse tissue. Upper row shows healthy control animal, bottom row shows tissue of NTS injected mouse on day 2, but not yet proteinuric. Scalebar – 5 µm. B – Serum creatinine of NTS induced mice. C – Serum urea of NTS induced mice. D – Podocyte damage score for proteomic data of laser dissected glomeruli of NTS injected mice. Data points: B+C – n=4 mice for Ctrl, NTS d2, NTS d3, NTS d5; n=3 mice for NTS d9, NTS d11. D – n=3 for all data points included. Statistical tests: ordinary one-way ANOVA with multiple comparisons for B+C. ns – not significant $p > 0.05$, * $p \leq 0.05$, ** $p \leq 0.01$, *** $p \leq 0.001$, **** $p \leq 0.0001$.

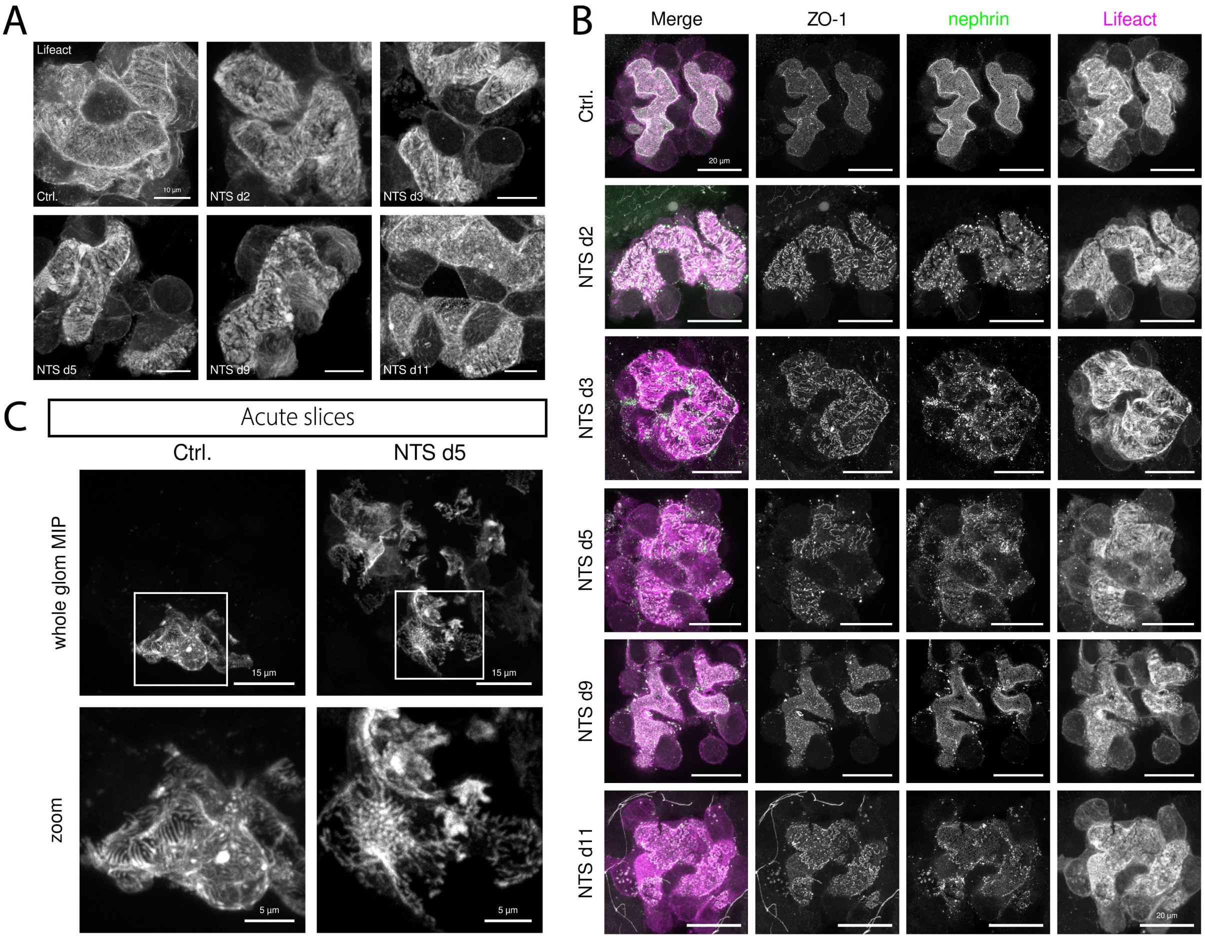


**Supplementary Figure 2: Lifeact.mScarlet-I staining reveals changes actin distribution during effacement.** A – Representative maximum intensity projection images of glomeruli stained for RFP/Lifeact of healthy control and NTS induced mice from day 2 to day 11. Scalebar – 10 µm. B – Maximum intensity projection images of endogenous Lifeact expression in living podocytes in AKS of iPod$^{Lifeact}$ mice. Left image column shows a healthy control, right column shows NTS d5. Zooms highlight Lifeact.mScarlet-I distribution of single podocytes. Scale bars: glomerulus – 15µm, zoom – 5µm. C – Representative maximum intensity projection images of glomeruli stained for nephrin (green), ZO-1 (white) and RFP/Lifeact (magenta) of NTS induced mice from day 2 to day 11. Scalebar – 20 µm.
